# Microbial Antioxidants Reduce ROS In Human Skin Cells Under Oxidative Stress

**DOI:** 10.64898/2026.01.15.699165

**Authors:** Shenghan Huang, Danhong Dong, Jinting Wan, Yang-Chi-Dung Lin, Hsien-Da Huang

**Author notes:** Correspondence: Hsien-Da Huang; Yang-Chi-Dung Lin.

## Abstract

Reactive oxygen species (ROS) play a dual role in cellular homeostasis, but excessive levels of ROS lead to oxidative stress, accelerating skin aging. Environmental stressors like UV radiation induce ROS overproduction, overwhelming endogenous antioxidant defenses and causing cellular damage. While the skin possesses an intrinsic antioxidant network that provides moderate protection, excessive oxidative stress can trigger inflammatory responses, thereby necessitating exogenous antioxidant intervention. Microbe-derived antioxidants (MA), produced via probiotic fermentation of sea buckthorn and chestnut rose, have shown promise in mitigating ROS-induced damage. In this study, we evaluated two MA formulations, MA1 and MA2, for their ability to scavenge free radicals and alleviate hydrogen peroxide (H_2_O_2_)-induced oxidative stress in human dermal fibroblasts (HDF) and dermal papilla cells (HDP). Both formulations displayed dose-dependent DPPH radical scavenging activity and enhanced cell viability at low concentrations. Under H_2_O_2_-induced oxidative stress, MA1 and MA2 effectively restored intracellular ROS to baseline levels, demonstrating significant cytoprotective effects. UHPLC–MS/MS profiling identified 12 compounds shared by both formulations, and Gene Ontology Biological Process enrichment analysis revealed that their associated target genes were significantly enriched in antioxidant-related pathways. Five compounds—adenosine, citric acid, 5-hydroxymethylfurfural, myricetin, and phenylalanine—emerged as key contributors to the observed antioxidative effects. Together, these findings highlight the potential of fermented microbial antioxidants to re-establish redox homeostasis in human skin cells and support their further development as therapeutic or cosmetic interventions targeting oxidative stress and skin aging. Given the heightened oxidative sensitivity of aged fibroblasts, MA’s ability to alleviate ROS may offer novel therapeutic strategies against skin aging and related pathologies.

## Introduction

Reactive oxygen species (ROS), predominantly generated in mitochondria [1], play a crucial role in maintaining cellular homeostasis, but excessive ROS can lead to oxidative stress, resulting in cellular damage and accelerated aging [2]. In skin tissues, particularly epidermal keratinocytes and dermal fibroblasts, ROS are frequently elevated following exposure to environmental stressors such as ultraviolet radiation (UVR) [3,4]. UVR induces immediate oxidative stress mediated by hydrogen peroxide (H_2_O_2_) [4–7]. UVR exposure also triggers the release of labile iron, which reacts with H_2_O_2_ and catalyzes the formation of highly reactive hydroxyl radicals via the Fenton reaction [4,7,8]. Excessive hydroxyl radical formation triggers oxidative damage across all major biomolecular classes — it induces strand breaks and base modifications in DNA, initiates lipid peroxidation of membrane polyunsaturated fatty acids, and oxidatively modifies amino acid residues in proteins [9,10]. When occurring in skin tissue, these events contribute to the hallmarks of premature aging, such as dermal collagen breakdown, loss of barrier function, and increased cellular senescence [11,12].

Skin possesses an intrinsic network of antioxidant systems, including enzymatic antioxidants such as glutathione peroxidase, superoxide dismutase, and catalase, as well as non-enzymatic antioxidants such as glutathione, ascorbic acid, and α-tocopherol, to maintain the ROS homeostasis [4]. However, these endogenous defenses can be overwhelmed by high oxidative stress conditions, such as prolonged or intense UVR exposure or acute inflammatory responses involving keratinocytes and infiltrating immune cells, resulting in substantial oxidative damage and accelerated skin aging [13].

Considering that chronic and acute oxidative stresses significantly impact cellular aging and integrity, external antioxidants have emerged as promising interventions to support the endogenous antioxidant defenses [4]. Microbe-derived antioxidants (MA) are produced from a mixture of chestnut rose (*Rosa roxburghii*) and sea buckthorn (*Hippophae rhamnoides*) through multi-stage solid–liquid complex fermentation by *Bacillus Subtilis, Lactobacillus, Clostridium Butyricum*, and *Saccharomyces cerevisiae*, [14,15]. Previous studies have shown that MA presents a novel and promising source of exogenous antioxidants with potential to effectively prevent ROS-induced cellular damage in various biological systems[14,16–20]. However, the comparative antioxidant efficacy of different MA formulations and their ability to regulate intracellular ROS levels in human skin–related cell types remain insufficiently characterized. Using an H_2_O_2_-induced oxidative stress model, this study evaluated the antioxidant effects of two MA formulations, MA1 and MA2, in human dermal papilla cells and dermal fibroblasts. Both MA1 and MA2 effectively restored intracellular ROS levels toward baseline. Dermal fibroblasts, particularly from older donors, exhibit heightened sensitivity to oxidative stress [21], evaluating the efficacy of MA in alleviating oxidative stress may provide critical insights to support their further development as functional food ingredients or nutraceutical components combating oxidative stress and skin aging.

## Materials and Methods

### DPPH Free Radical Scavenging Capacity Assay

Stock solutions of MA1 and MA2 were provided by Jianghan Biotechnology (Shanghai) Co., LTD. The free radical scavenging capacity of MA1 and MA2 were measured with DPPH Free Radical Scavenging Capacity Assay Kit (Sangon Biotech, NO. D799008-0100) according to the manufacturer’s protocol. Briefly, stock solutions of MA1 and MA2 were serially diluted to achieve final concentrations of 5%, 2.5%, 1.25%, 0.625%, and 0.3125% (v/v). A working solution of DPPH (190 µL) was then added to each sample well, and the microplate was incubated in the dark at 21°C for 30 minutes. The optical density (OD) of each well was measured at 515 nm using a microplate reader (EPOCH2, BioTek, USA). The blank control was prepared with Reagent 1 instead of the DPPH working Solution. Vitamin C was used as the positive control. The radical scavenging rate was calculated by the following formula:

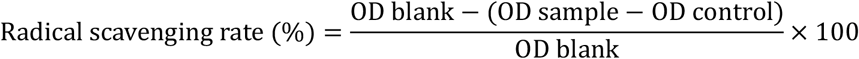

### Culture of Human Dermal cells

Primary human dermal papilla cells (HDP) (Delf-10698) and Primary human dermal fibroblasts (HDF) (Delf-16276) cells were obtained from Delf, Hefei Wanwu Biotechnology Co., LTD (Hefei, China). HDP and HDF cells were maintained in Primary Fibroblast Culture System (Delf, No. Delf-15764). The cells were cultured at 37°C with 5% CO_2_ and passaged after reaching 80% confluency for no more than 10 passages.

### Cell Treatments

For MA1, MA2, and H_2_O_2_ treatments, solutions were diluted in culture media to the final concentrations indicated in each experiment. For H_2_O_2_ induced oxidative stress, the cells were treated with 400 µM H_2_O_2_ (aladdin, No. H112515) for 30 minutes. Then, H_2_O_2_-containing media was replaced with either MA1, MA2 or culture media. The cells were incubated for 48 hours prior to subsequent analysis.

### Cell Viability Assay

The cytotoxicity of MA1 and MA2 was assessed with Enhanced CCK-8 Kit (Beyotime, No. C0046) according to the manufacturer’s protocol. In brief, cells were cultured in 96-well culture plates for 24 hours before being treated with various concentrations of MA1 or MA2 diluted in culture media for 48 hours. The old culture media were completely removed, and cells were treated with CCK-8 for 1 hour, and the absorbance at 450 nm was subsequently measured using a microplate reader (EPOCH 2, BioTek, USA).

### Reactive Oxygen Species Assay

The intracellular levels of ROS were detected with 2, 7-Dichlorofuorescin Diacetate (DCFH-DA) (NJJCBIO, No. E004-1-1) according to the manufacturer’s protocol. In brief, cells were seeded in 6-well culture plates for 24 hours before treatments. The fluorescence intensity of 5000 cells was determined with a flow cytometer (CytoFLEX S, Beckman Coulter), and the data was analysed in CytExpert 2.4.

### Compound Detection with UHPLC-MS/MS

Samples were prepared with 50% methanol and 50% of each MA, sonicated for 5 minutes, and centrifuged at 20,000 g for 10 minutes. Subsequently, 8 µL of the supernatant was injected in both positive and negative ion modes for UHPLC-MS analysis. The UHPLC model used was LC-30A (SHIMADZU, JAPAN). The column used was Shim-pack GIST, dimension 2.1×100mm (HSS) (SHIMADZU, Japan). The water phase used contained 99.9% water, and 0.1% formic acid. The organic phase used contained 100% acetonitrile. Gradient elution was conducted with the following program: 0-2 min, from 0% to 5% B; 2-22 min, 99% B; 22-26 min, 99% B; 26-30 min, from 99% to 5% B. Flow rate: 0.3 mL/min; column temperature: 30°C; injection volume: 10 μL. Spectrometry analysis was performed using Sciex TripleTOF 6600+ system with the following parameters: curtain gas (CUR) set to 35 psi, gas source 1 (GS1) at 55 psi, gas source 2 (GS2) at 55 psi, ion spray voltage (ISVF) at 5500 V/-4500 V (positive/negative), and ion source temperature (TEM) at 550°C. Compound identification was performed using SCIEX OS software with the TCM MS/MS Library 2.2. Only compounds with a library score greater than 95 were considered confidently identified. The identified compounds in MA1 and MA2 are stored on Zenodo at: https://doi.org/10.5281/zenodo.18263939.

### Gene Ontology Biological Process Enrichment Analysis

The compounds shared by both MA1 and MA2 were used to retrieve their corresponding human protein targets from the DrugBank databases [22]. The gene symbols associated with these targets, as provided directly by DrugBank, were used for Gene Ontology Biological Process (GO BP) enrichment analysis using the clusterProfiler package in R. GO BP terms with a p-value < 0.05 and q-value < 0.05 were considered significantly enriched.

### Statistical Analysis

Statistical analyses were performed using GraphPad Prism 8.0.2 software (San Diego, CA, USA). One-way analysis of variance (ANOVA) followed by Dunnett’s post hoc test was used for multiple group comparisons. Data are presented as mean ± standard error of the mean (SEM).

## Results

A schematic overview of the experimental framework to identify the intracellular ROS effect of MA treatment is presented in Figure 1, where HDF and HDP cells were treated with different concentrations of MA1 or MA2 to obtain optimal administration concentrations.

**Figure 1.**
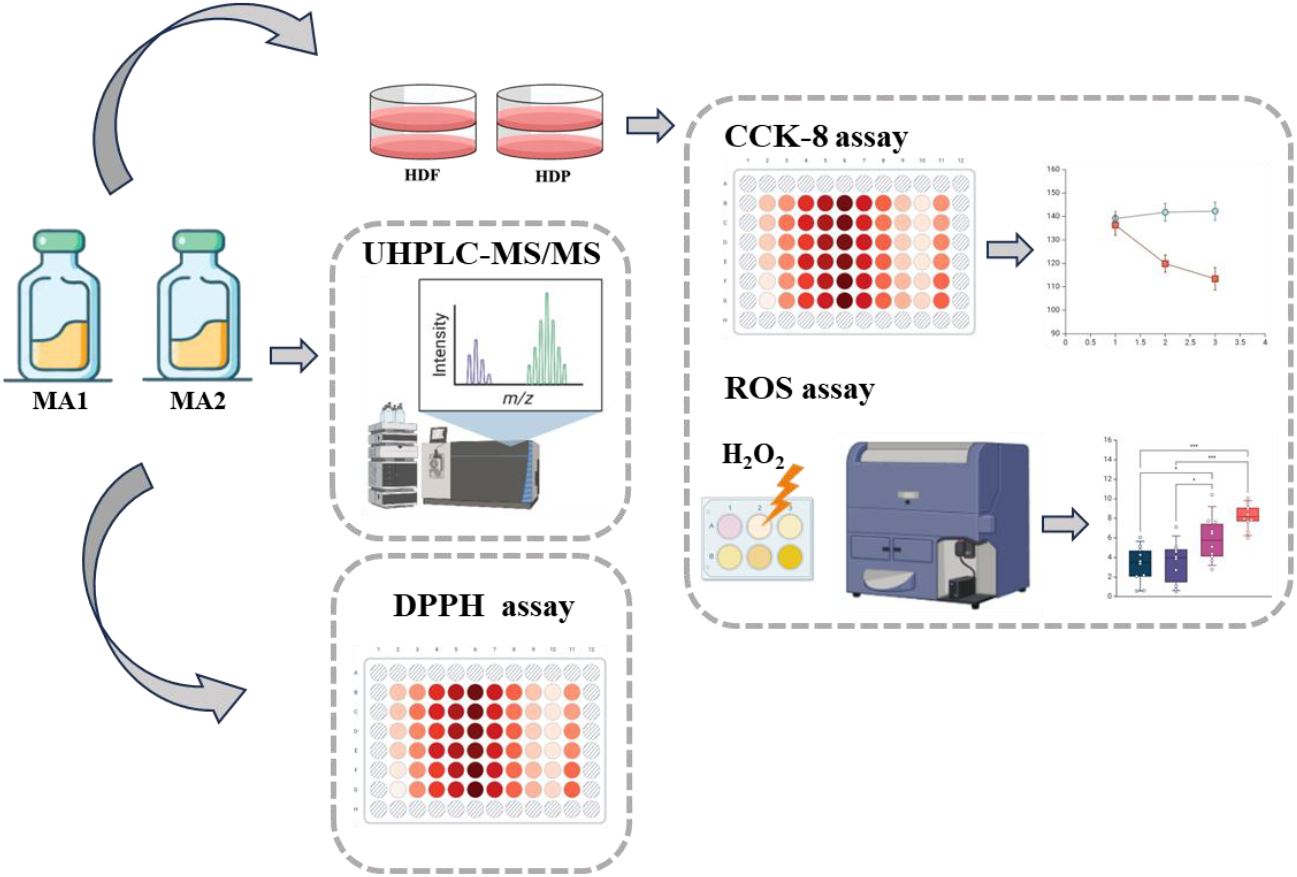
Experimental design. The free radical scavenging capacities of MA1 and MA2 were evaluated using the DPPH assay. The effects of MA1 and MA2 on HDF and HDP cells were assessed with the CCK-8 assay, while intracellular ROS levels were determined by flow cytometry. The chemical compositions of MA1 and MA2 were identified by UHPLC–MS/MS analysis.

### MA1 and MA2 Display Dose-Dependent Free Radical Scavenging Activity

Both MA1 and MA2 displayed dose-dependent radical scavenging activity (Figure 2). The antioxidant potential of MA1 and MA2 was evaluated using the DPPH radical scavenging assay. The EC50 value of vitamin C, MA1, and MA2 was determined to be 0.13 mg/µL for vitamin C, 2.523% (v/v) for MA1, and 4.681% (v/v) for MA2.

**Figure 2.**
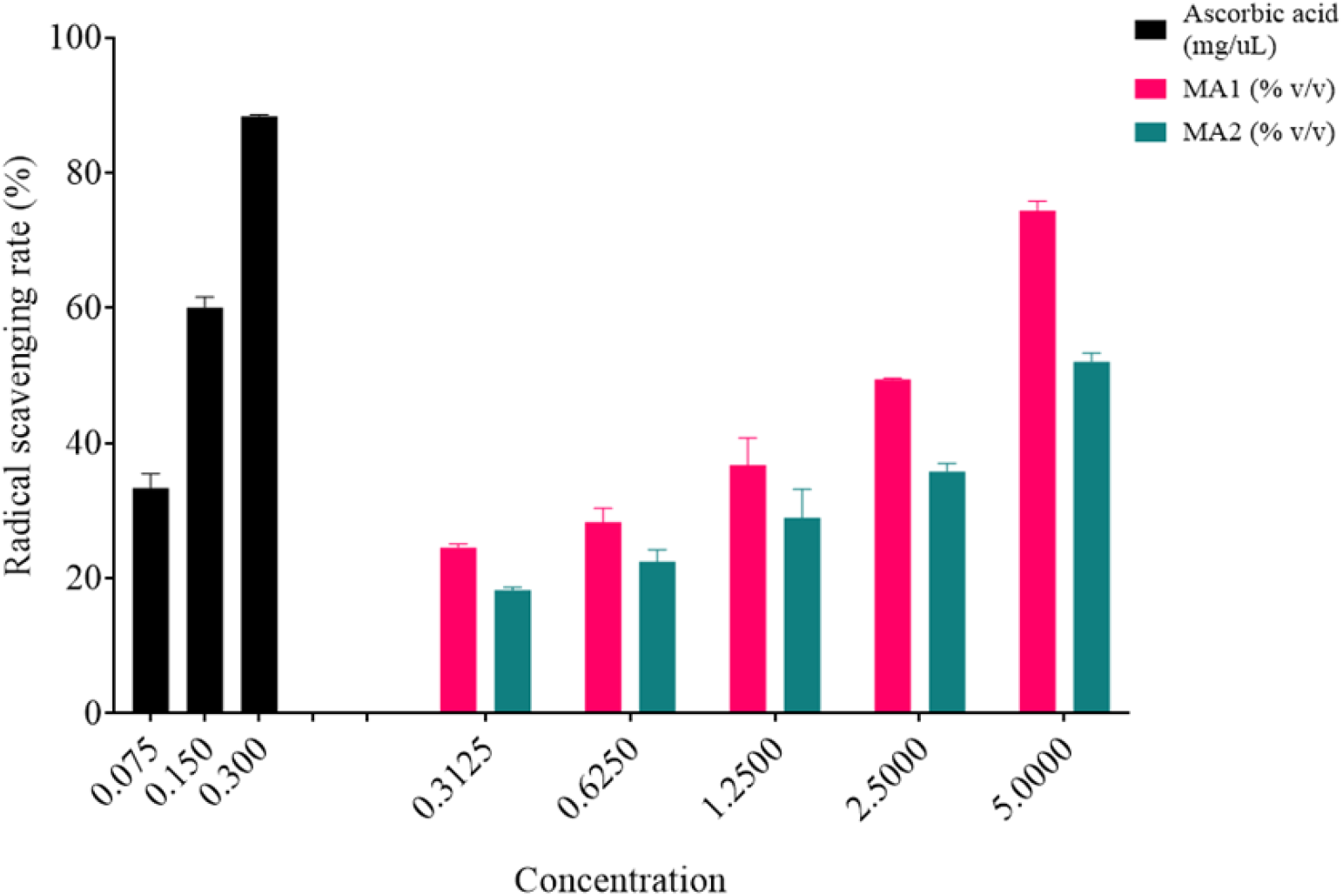
DPPH radical scavenging activity of MA1 and MA2. The data are presented as the means ± SEMs of 2 independent experiments.

### MA1 And MA2 Restore H_2_O_2_-Induced ROS Levels in HDF And HDP Cells

The effects of MA1 and MA2 on the viability of HDF and HDP cells were evaluated using the CCK-8 cytotoxicity assay (Figure 3a). The untreated cells served as the control. Cells treated with MA1 or MA2 at concentrations below 2% (v/v) showed increased viability compared with the control. For MA1, the highest viability was observed at 0.62% (v/v) in HDF cells and 1% (v/v) in HDP cells, whereas for MA2, peak viability occurred at 1.25% (v/v) in HDF cells and 1% (v/v) in HDP cells. Both HDF and HDP cells retained normal morphology when treated with MA1 or MA2 at their respective peak viability concentrations (Figure 3b). These peak concentrations were used for subsequent experiments. The effects of MA1 and MA2 on intracellular ROS levels were assessed using the DCFH-DA assay (Figure 3c). Under normal culture conditions, neither MA1 nor MA2 significantly altered intracellular ROS levels. Cells exposed to 400 µM H_2_O_2_ showed a significant increase in ROS levels, while treatment with either MA1 or MA2 reduced intracellular ROS in H_2_O_2_-stimulated cells to levels not significantly different from the control.

**Figure 3.**
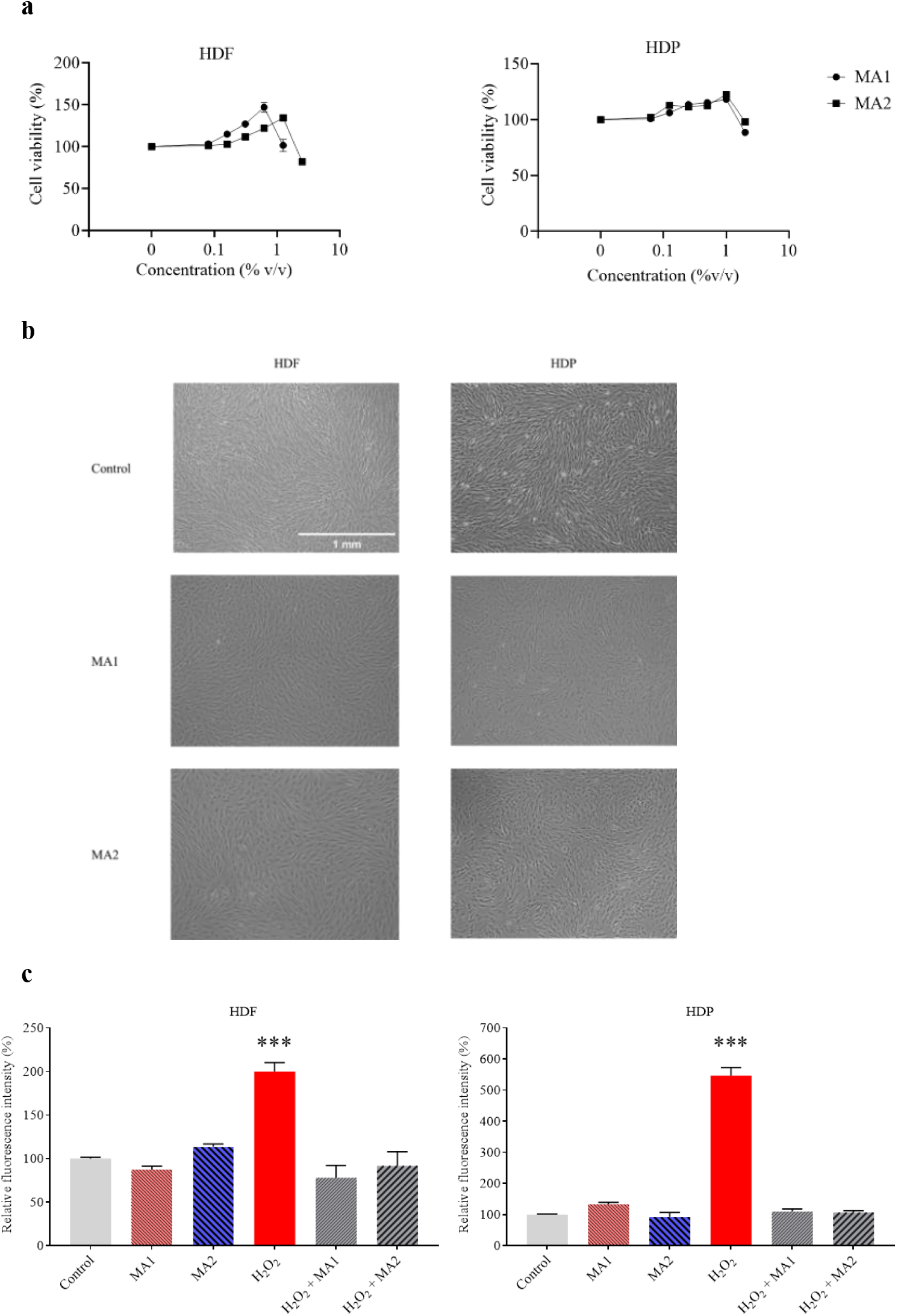
Effect of MA1 and MA2 on HDF and HDP cells. (a) Cell viability of HDF and HDP treated with MA1 or MA2. (b) Stereomicroscopic images of HDF and HDP treated with MA1 or MA2 for 48 hours. HDF was treated with 0.625% (v/v) MA1 or 1.25% (v/v) MA2, and HDP was treated with 1% (v/v) MA1 or MA2. (c) Effect of MA1 or MA2 on the intracellular ROS levels of HDF or HDP cells under normal culturing conditions, and under 400 µM H_2_O_2_-induced oxidative stress. For HDF cells MA1 at 0.625% (v/v) and MA2 at 1.25% (v/v) were used. For HDP cells 1% (v/v) of MA1 or MA2 were used. The data are presented as the means ± SEMs of 3 independent experiments. The data were statistically evaluated by one-way ANOVA followed by Dunnett’s multiple comparisons test to compare each treatment group to the control; * p < 0.05, **< 0.005, and ****< 0.0001 compared to the control.

### Common Compounds In MA1 And MA2 Are Associated With Enrichment In Antioxidant-Related Biological Processes

UHPLC–MS/MS was conducted to characterize the compounds contributing to the ROS-lowering effects shared by MA1 and MA2. A total of 36 compounds were identified in MA1, and 18 compounds were found in MA2. Interestingly, 12 compounds were common to both MA1 and MA2 (Table 1). To understand the biological implications of these common compounds, Gene Ontology Biological Process (GO BP) enrichment analysis was conducted. This analysis revealed that these 12 common compounds are significantly enriched in GO BP related to anti-oxidation (Figure 4). The enrichment in anti-oxidation related pathways strongly supports the experimental findings regarding the ROS scavenging capabilities of MA1 and MA2. Based on the GO enrichment analysis, we propose that MA1 and MA2 act through adenosine, citric acid, hydroxymethylfurfural, myricetin, and phenylalanine to restore H_2_O_2_-induced ROS levels (Figure 5).

**Table 1.** Common compounds identified in MA1 and MA2 and their respective target genes.

| Library Hit | Retention Time (min) | m/z | Target Genes |
| --- | --- | --- | --- |
| 4-Hydroxybenzoic acid | 6.28 | 137.0249 | Amine oxidase [flavin-containing] B (MAOB) |
| 5-Hydroxymethylfurfural | 3.06 | 127.0379 | Hemoglobin subunit alpha (HBA1) |
| Adenosine | 1.75 | 268.1021 | Adenosine receptor A1 (ADORA1), Adenosine receptor A2a (ADORA2A), Adenosine receptor A2b (ADORA2B), Adenosine receptor A3 (ADORA3) |
| Betaine | 28.17 | 118.0858 | Betaine--homocysteine S-methyltransferase 1 (BHMT) |
| Citric acid | 1.02 | 191.0542 | Aldo-keto reductase family 1 member B1 (AKR1B1), Transient receptor potential cation channel subfamily V member 4 (TRPV4), Citrate synthase, mitochondrial (CS), Proto-oncogene tyrosine-protein kinase Src (SRC), Ribonuclease pancreatic (RNASE1), Eosinophil cationic protein (RNASE3), L-amino-acid oxidase (IL4I1), Carboxy-terminal domain RNA polymerase II polypeptide A small phosphatase 1 (CTDSP1), Hepatocyte growth factor-regulated tyrosine kinase substrate (HGS), Malate dehydrogenase, mitochondrial (MDH2), Carboxypeptidase B (CPB1), U6 snRNA-associated Sm-like protein LSM6 (LSM6), Pleckstrin homology domain-containing family A member 1 (PLEKHA1), Complement component C8 gamma chain (C8G)<br>Adenine phosphoribosyltransferase (APRT), Inosine triphosphate pyrophosphatase (ITPA) |
| Dihydromyricetin | 7.70 | 321.0578 | N/A |
| Myricetin | 0.77 | 317.0544 | Phosphatidylinositol 4,5-bisphosphate 3-kinase catalytic subunit gamma isoform (PIK3CG), Tyrosine-protein kinase JAK1 (JAK1) |
| Myricitrin | 8.67 | 463.0877 | N/A |
| Nicotinic acid | 1.52 | 124.0384 | Hydroxycarboxylic acid receptor 3 (HCAR3), Hydroxycarboxylic acid receptor 2 (HCAR2), Nicotinate-nucleotide pyrophosphorylase [carboxylating] (QPRT), Nicotinamide N-methyltransferase (NNMT), Diacylglycerol O-acyltransferase 2 (DGAT2) |
| Phenylalanine | 7.32 | 415.1032 | Tyrosine 3-monooxygenase (TH), Tyrosine aminotransferase (TAT), Large neutral amino acids transporter small subunit 2 (SLC7A8), Phenylalanine--tRNA ligase alpha subunit (FARSA)<br>Phenylalanine-4-hydroxylase (PAH), Phenylalanine--tRNA ligase, mitochondrial (FARS2), Phenylalanine--tRNA ligase beta subunit (FARSB) |
| Puerarin | 0.92 | 138.0545 | 5-hydroxytryptamine receptor 2C (HTR2C) |
| Trigonelline | 2.57 | 166.0857 | N/A |

**Figure 4.**
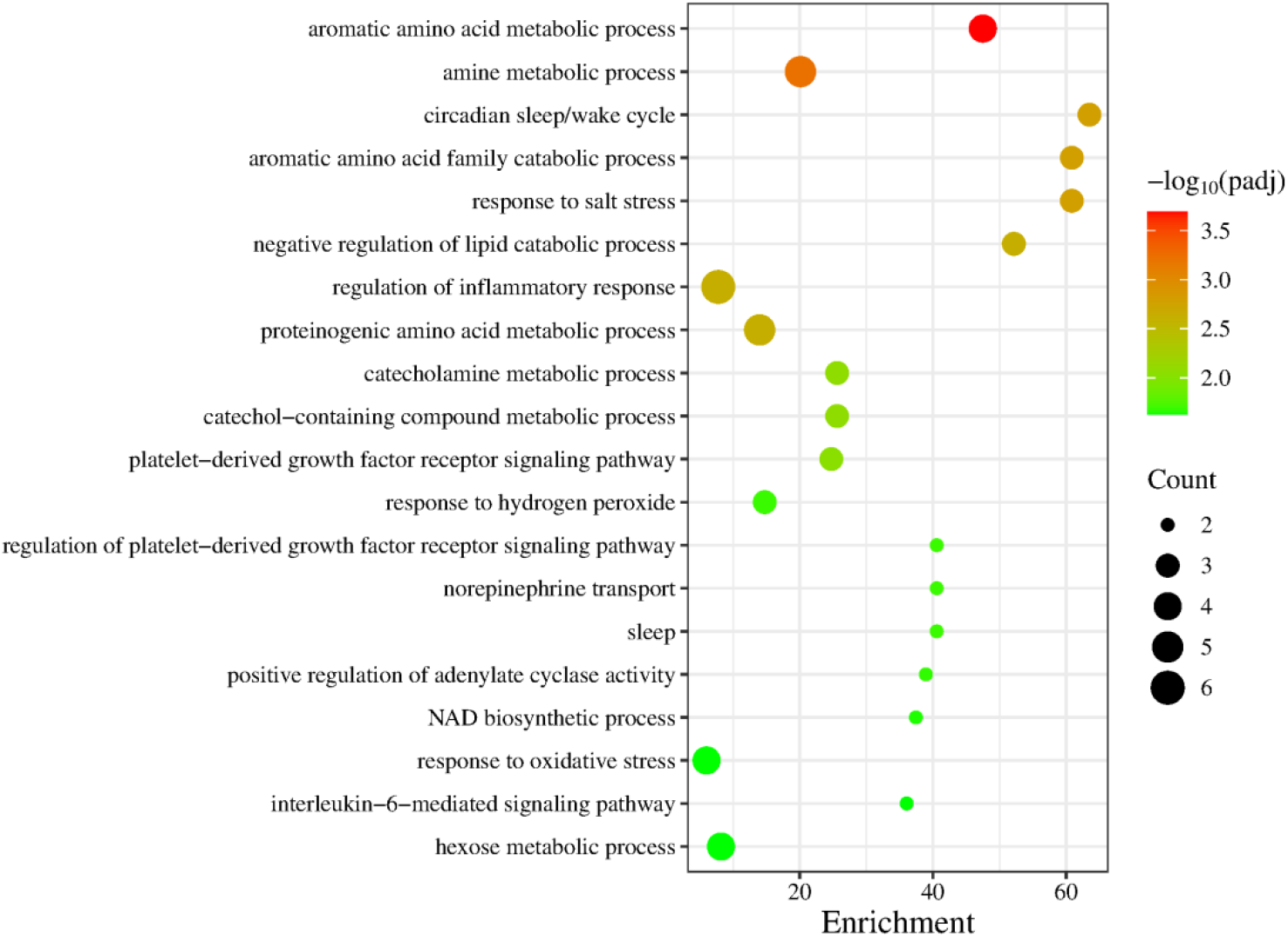
GO Biological Process enrichment analysis of the target genes of the common compounds between MA1 and MA2.

**Figure 5.**
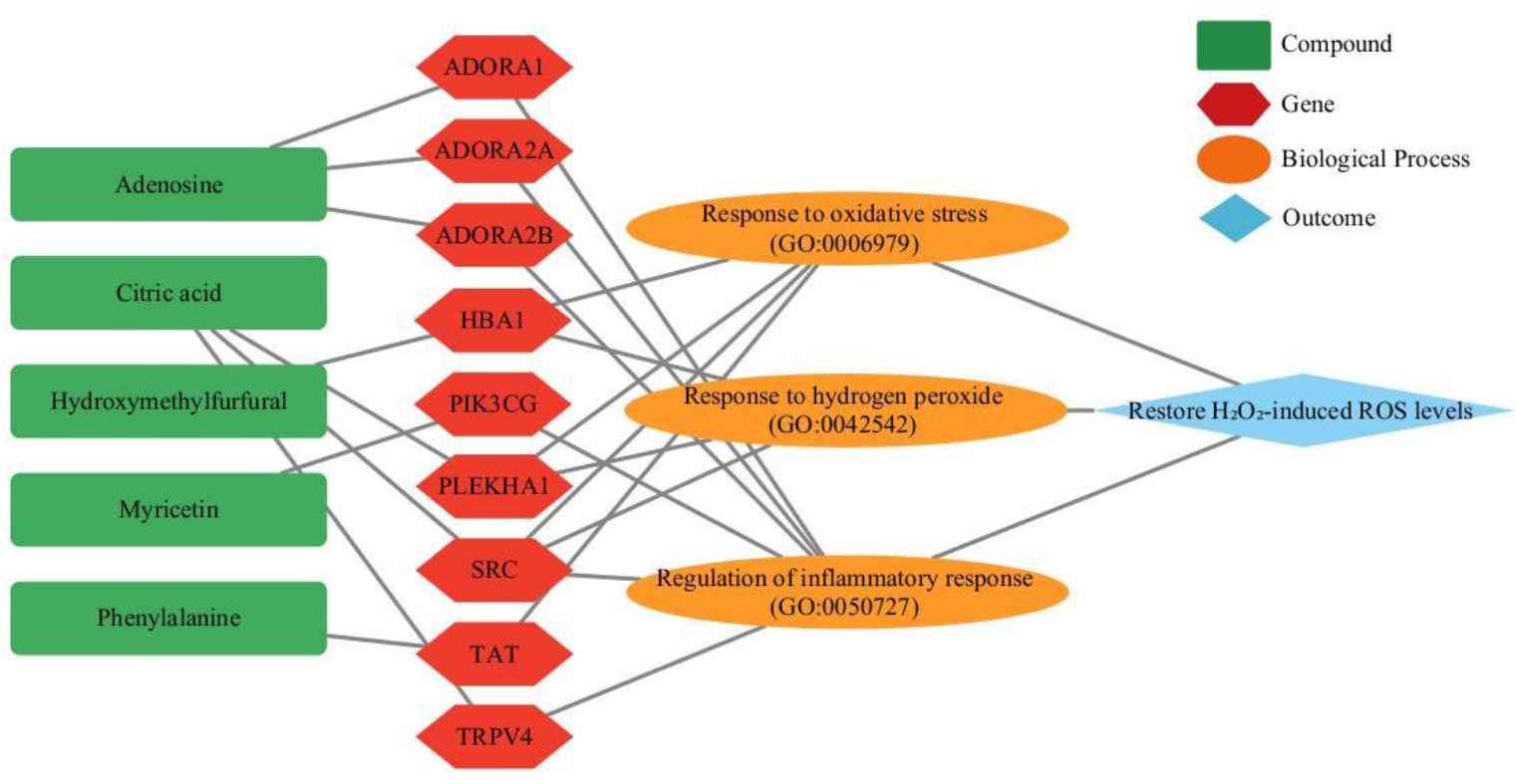
Proposed antioxidation mechanism of MA1 and MA2.

## Discussion

This study demonstrates that MA, produced through the fermentation of sea buckthorn and chestnut rose, effectively mitigates oxidative stress in human dermal fibroblasts and dermal papilla cells. Our findings reveal that two distinct MA formulations, MA1 and MA2, exhibit significant dose-dependent free radical scavenging activity and can restore intracellular ROS levels to a normal baseline after being elevated by H_2_O_2_-induced oxidative stress. These results align with a growing body of research highlighting the potential of fermented natural products as a source of potent antioxidants [23,24].

The observed antioxidant capacity of MA1 and MA2 is attributed to a complex interplay of bioactive compounds generated during the fermentation process. Our UHPLC-MS/MS analysis identified 12 common compounds in both formulations. GO BP enrichment analysis of the target genes for these compounds revealed a significant enrichment in antioxidant-related processes. These processes converge on five key compounds—adenosine [25–28], citric acid [29,30], 5-hydroxymethylfurfural [31,32], myricetin [33–36], and phenylalanine [37,38]—all of which have been reported to possess antioxidant properties, providing a molecular basis for the observed ROS-regulating effects. This is consistent with previous studies that have identified these and similar compounds in fermented plant extracts and have demonstrated their antioxidant properties [39,40]. We propose that the synergistic formulation of these five key compounds plays a pivotal role in re-establishing intracellular ROS equilibrium under H_2_O_2_–induced oxidative stress.

The protective effect of MA against oxidative stress is particularly relevant in the context of skin aging. Dermal fibroblasts, especially from older donors, are known to be more susceptible to oxidative damage, which contributes to the visible signs of aging, such as wrinkles and loss of elasticity [21]. By effectively reducing ROS levels in these cells, MA may help to preserve cellular function and integrity, thereby slowing down the aging process. The capacity of MA to reestablish ROS homeostasis under H_2_O_2_-induced oxidative conditions implies potential therapeutic applications in protecting the skin from environmental aggressors, such as UV radiation, which are recognized as major sources of hydroxyl radical generation [5].

While this study provides compelling evidence for the antioxidant potential of MA under controlled in vitro conditions, several limitations should be acknowledged. The present work is limited to in vitro models, and direct validation of MA1 and MA2 in more complex biological systems, such as in vivo animal models, is warranted. Notably, previous studies have demonstrated that MA effectively reduces intestinal oxidative stress in weanling piglets [14], alleviates high-fat diet–induced lipid dysregulation and metabolic inflammation in the liver and adipose tissue of mice [18], and protects IPEC-1 cells from H_2_O_2_-induced oxidative stress, inflammation, and tight junction protein disruption through activation of the Nrf2 pathway and subsequent inhibition of the ROS/NLRP3/IL-1β signaling axis [20]. These findings support the physiological relevance of MA but do not substitute for in vivo validation of the specific formulations examined in this study. Additionally, while we have identified several common compounds in MA1 and MA2, the synergistic or antagonistic interactions between these compounds remain to be elucidated.

In conclusion, this study provides strong evidence that microbe-derived antioxidants from fermented sea buckthorn and chestnut rose can effectively protect human dermal cells from excessive oxidative stress. Collectively, these findings support the potential use of these natural formulations as functional foods to combat skin aging and other oxidative stress-related pathologies. Further research is warranted to fully elucidate the molecular mechanisms underlying the antioxidant effects of MA and to evaluate their efficacy and safety.

